# Simpatico: accurate and ultra-fast virtual drug screening with atomic embeddings

**DOI:** 10.1101/2025.06.08.658499

**Authors:** Jeremiah Gaiser, Travis J. Wheeler

**Affiliations:** School of Information, 85721, Arizona, USA; College of Pharmacy, 85721, Arizona, USA

**Keywords:** Drug screening, Representation learning, Embeddings, Protein drug interactions

## Abstract

Virtual screening, the in-silico assessment of large libraries of small molecules for binding to a therapeutic protein target, is a critical early step in drug discovery. The dominant approach, molecular docking, requires a separate calculation for each protein-molecule pair, and is too slow to apply alone at the billion-compound scale of modern compound libraries. A recent embedding-retrieval paradigm, exemplified by DrugCLIP, addresses this bottleneck by training deep models to map proteins and small molecules into a shared embedding space, such that proteins are co-located with their likely binding partners; candidate ligands can then be retrieved directly by nearest-neighbors search, with no per-pair calculation. However, current embedding-retrieval methods collapse each protein and each ligand into a single embedding, creating an information bottleneck that limits the representation of partial or alternative binding compatibility. We present `simpatico`, an embedding-retrieval virtual screening tool that instead produces a unique embedding for each atom in a protein pocket or ligand. A CLIP-style contrastive objective trains these atomic embeddings so that protein-ligand atom pairs known to interact are nearby in embedding space. To screen a protein target, each protein-atom embedding is used as a query against a vector database of precomputed small-molecule atomic embeddings, returning the closest atoms in the library; a simple aggregation step assigns a binding score to each candidate molecule containing retrieved atoms. Where prior retrieval-based methods index one vector per ligand, simpatico indexes one per heavy atom; query time grows sublinearly in library size.

On challenging decoy benchmarks, `simpatico` achieves state-of-the-art predictive accuracy, outperforming recent dense-retrieval methods despite training on only ∼15,000 protein-ligand complexes from PDBBind, with no pretraining and no 3D ligand pose estimation. `Simpatico` also exceeds the accuracy of physics-based docking and deep-learning-augmented docking methods, is competitive with diffusion-based docking, and runs orders of magnitude faster than all three. `Simpatico` is open source software; all code, weights, and data may be accessed at https://github.com/TravisWheelerLab/Simpatico.

## Introduction

Virtual screening (VS [23])) is the in-silico assessment of large compound libraries to identify candidates likely to bind a therapeutic target, allowing investigators to narrow the search to a tractable number of compound families before any wet-lab work. The scale of libraries used for VS has grown dramatically; “large” now routinely means billions of compounds [9, 25, 27]. It has been observed [16] that top-scoring molecules from ultra-large-library docking campaigns, including experimentally confirmed actives, tend to occupy regions of chemical space distinct from bio-like molecules, a pattern that does not emerge at smaller library scales. Methods unable to screen at billion-scale forfeit access to that chemistry.

Among the most popular methods employed by VS is molecular docking. Classical docking tools such as AutoDock Vina [8], Glide [11], and GOLD [13] take protein-pocket and small-molecule structures as input and perform a principled search over the probable ways that the hypothetical ligand would rest in the protein pocket. These resting conformations can then serve as the input for energy-based scoring functions to produce an estimate of protein-ligand binding affinity.

Docking and docking-based scoring have recently been augmented in two ways by deep learning. First, learned scoring functions (e.g., Gnina [17], RF-Score [29]) replace or supplement classical energy terms with neural network–based estimates of binding affinity. Second, generative deep-learning models replace classical pose enumeration entirely. Diffusion-based docking methods such as DiffDock [6] and SurfDock [5] iteratively denoise candidate ligand poses, and related diffusion machinery underlies general biomolecular structure-prediction models (such as AlphaFold3 [1] and Boltz-2 [20]) that can predict full protein-ligand complexes. Other GNN-based approaches such as KarmaDock [30] instead predict bound conformations directly in a single forward pass. However, all these docking approaches still require performing a new calculation for each pose of each protein-molecule pair, and each calculation takes seconds or minutes to complete. In practice, this precludes the use of docking alone to screen large databases of billions of molecules.

This bottleneck has motivated development of a dense-retrieval approach to VS, in which models map proteins and small molecules into a shared high-dimensional embedding space such that proteins are co-located with their binding partners. A protein query then retrieves candidate ligands by nearest-neighbors search over a precomputed vector database (e.g., FAISS [7]), bypassing per-pair calculation entirely. The recent dense-retrieval method DrugCLIP [12] established the viability of this paradigm, demonstrating that embedding retrieval can identify binding candidates more effectively than docking, with much greater speed. Using GNN encoders trained with a CLIP-style contrastive objective [21], DrugCLIP embeds protein pockets and small molecules into a shared space, such that pocket embeddings sit near the embeddings of their known active ligands. However, collapsing each protein pocket and ligand into a single embedding creates an information bottleneck: the many possible interaction modes available to each protein and ligand must be compressed into a single location in embedding space. This limits the embeddings’ ability to express nuanced relationships, such as when a ligand binds to multiple proteins through different sets of interactions, or when significant portions of the ligand do not associate strongly with the protein.

We present our retrieval-based VS tool, `simpatico` (Simple atomic-interaction prediction with Contrastive learning). Like DrugCLIP, `simpatico` encodes proteins and small molecules in a dual graph neural network architecture trained under a CLIP-style loss function. However, our model differs in a key aspect: rather than reduce each protein and each small molecule to a single fixed-dimensional vector, `simpatico` outputs a unique vector for each atom, and trains these embeddings so that protein-ligand atom pairs known to interact are nearby in embedding space. This enables a powerful variant of the dense-retrieval search task: vector representations of individual protein atoms are used as queries to probe a database of small-molecule atoms for physicochemically complementary partners.

For each candidate molecule with at least one retrieved atom, we aggregate hits into a binding score. Under the simple intuition that true binders should accumulate more physicochemical complements than non-binders, we sum the embedding-distance-weighted contributions of each returned atom belonging to that molecule.

In a mock screen for actives in a large database of random molecules (∼100M), `simpatico` demonstrates high enrichment at very small sampling ratios, with some targets achieving enrichment values of several thousand fold, while requiring only ∼ 14 seconds per million small molecules for a typical protein pocket target.

When applied to challenging decoy screening tasks, `simpatico` achieves state-of-the-art predictive accuracy (enrichment factor). It outperforms recent dense-retrieval based screening approaches while requiring zero pretraining, no 3D ligand pose estimation, and *significantly* less training data (in some cases, only ∼ 15,000 PDBBind complexes, after filtering homologs of test-set targets). `simpatico` also shows greater predictive accuracy than physics-based docking (AutoDock Vina[8]) and Deep Learning–augmented docking (Gnina [17]), and is competitive with diffusion-based docking (SurfDock [5]), while running orders of magnitude faster than all three.

`Simpatico` is open source software; all code, weights, and data are available at https://github.com/TravisWheelerLab/Simpatico.

### Graph Neural Networks for molecules

Graph neural networks (GNNs; see [24] for a practical introduction) are deep learning architectures designed to operate directly on graph-structured data, learning to update node features iteratively through multiple message-passing layers. This design allows GNNs to model data with spatial and non-Euclidean relationships, making them ideally suited for molecular structures. They naturally represent chemical structures through a node-edge framework, wherein atoms correspond to nodes and chemical bonds correspond to edges.

Formally, a GNN operates on a graph *G* = (*V, E*), defined by a set of nodes *V* and a set of edges *E*. A GNN updates node features iteratively through multiple message-passing layers. Specifically, at layer *l*, each node *v* ∈ *V* is updated based on information aggregated from its local neighborhood. For each node *v*, let *N*(v) ⊆ *V* denote the set of neighbor nodes connected to *v*, and let *e*_*vn*_ represent edge features associated with the edge connecting nodes *v* and *n* ∈ *N*(*v*). The update rule at layer *l* for node *v* can be formally defined as follows:

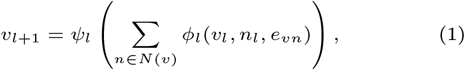

where *ϕ*_*l*_ and *ψ*_*l*_ are learnable nonlinear functions. Specifically, *ϕ*_*l*_ processes and aggregates information from each neighboring node and associated edge features, and *ψ*_*l*_ generates an updated node representation based on the aggregated neighborhood information.

The parameters *θ* = {*ϕ, ψ*} are optimized by minimizing a suitable loss function *L*(*V*_*Z*_), computed using the node representations *V*_*Z*_ obtained from the final layer *Z*.

### Contrastive Learning

Contrastive learning is a self-supervised learning technique widely used in representation learning. The basic strategy of contrastive learning is to update network parameters in a way that pulls representations closer together in embedding space if their original inputs belong to the same underlying class and pushes them apart if their original classes differ. Over many training batches, the network learns a structured embedding space in which underlying classes can be distinguished. To achieve this, the network must learn a rich set of features corresponding to specific classes.

### Notation

For clarity, we introduce the following notation. A protein-ligand complex comprises a protein pocket and a small molecular compound (ligand). When it is necessary to distinguish unique pockets or compounds, we employ subscripts explicitly denoting their distinct identities. When discussing a collection of pockets ℙ, we may refer to an individual pocket as *P*_*a*_. Similarly, each ligand *M*_*b*_ belongs to a collection of small molecules 𝕄. Proteins and ligands themselves consist of atoms, with *p*_*a,i*_ representing the *i*-th atom in the *a*-th pocket from the set ℙ, and *m*_*b,j*_ representing the *j*-th atom in the *b*-th ligand from the set 𝕄. For notational convenience, when pocket/ligand identity is clear, we remove the pocket/ligand identifier, leaving *P, M, p*_*i*_, and *m*_*j*_.

A small molecule *M* might be known to associate with a specific pocket *P*. This relationship is denoted as *M* ∼ *P*, or equivalently, *P* ∼ *M*. If we know that small molecule *M* explicitly does *not* associate with protein pocket *P*, we write that *M* ≁ *P*.

For any pocket *P* and some known active *M* such that *P* ∼ *M*, the binding affinity between the two chemical structures will depend on the interactions between their atoms. We consider any two atoms *p*_*i*_ and *m*_*j*_ belonging to a protein *P* and compound *M* to be *interacting* if they are observed within 4 angstroms (Å) of one another in a resolved crystal structure of the complex *PM*. In shorthand, we express this interaction as *p*_*i*_ ∼ *m*_*j*_ (or equivalently *m*_*j*_ ∼ *p*_*i*_).

The above notation is useful for describing physical relationships among protein pockets, small molecular compounds, and their constituent atoms. The aim of `simpatico` is to train an encoding function *f*(·|*θ*) to project these atoms into a high-dimensional embedding space ℝ^*d*^. To differentiate an atom *p*_*i*_ or *m*_*j*_ from its embedding representation, we denote atom embeddings as 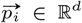, such that 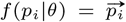 (and similarly, 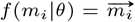). The notation for describing the set of *k* nearest neighbors for some vector 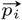 in high dimensional space is 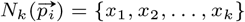, where each *x*_*l*_ is one of the *k* closest points to 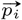 according to a specified metric (in this case, the Euclidean (*L*_2_) distance). If an embedding 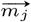 belongs to the set of *k* nearest neighbors 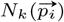, we write 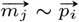.

When we discuss the collection of atomic embeddings of all the atoms in some pocket *P* or compound *M*, we will refer to 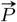 or 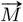. To round out our notation, we will refer to a collection of 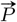 or 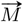 with 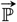 or 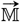, respectively.

## Methods

### Motivation

The `simpatico` method is based on the concept of a desirable semantic space for atomic embeddings of proteins and small molecules, defined by the characteristic that if a small molecular compound atom *m*_*j*_ and a protein atom *p*_*i*_ from a resolved protein-ligand complex interact (are neighbors) in 3D space (*m*_*j*_ ∼ *p*_*i*_), then they should also be neighbors in the learned embedding space 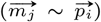. With a trained model, atomic representations in this space are organized according to some implicit notion of physicochemical complementarity, or the likelihood that they interact with one another. Note that the probability of noncovalent bonding between any two atoms is not simply a function of their species, but is in fact a geometrically complex collection of variables including the species and spatial composition of neighboring atoms; this means that the model must learn a representation rich enough to integrate local structural context, not merely encode atom type.

Per-atom embeddings allow each atom to be matched independently against a precomputed database, so even a single strongly complementary atom can retrieve its parent molecule. This capability is inaccessible to whole-molecule embeddings.

### Data

Evaluation was performed using standard benchmarks: DEKOIS [2], DUD-E [18], and LIT-PCBA [26] (see Supplementary Materials for descriptions of these data sets). To evaluate high-throughput speed and enrichment, these were supplemented by compounds from the Enamine REAL database [9].

Training data were taken from PDBBind [15], a curated dataset of 19,443 high quality protein-ligand complexes spanning 3,876 distinct proteins, sourced from the Protein Data Bank [4]. PDBBind entries provide simplified (no ions, water or extraneous heteroatoms) structures of ligands in complex with their protein target.

For each evaluation set, MMseqs2 was used to identify any proteins in PDBBind with ≥ 90% sequence identity to any proteins in the test data. Any training samples involving these proteins were withheld from the training data for evaluation on that benchmark. This resulted in a significant reduction of available training data. This filtering removed 4,172 training samples for DUD-E (yielding 15,271 remaining, a >20% reduction), 2,703 for DEKOIS, and 237 for LIT-PCBA, which has fewer targets and thus fewer homologs.

### Compound Graph Construction

Small molecule compounds (drugs) are converted to Pytorch Geometric (PyG)-formatted graphs [10]. Each heavy atom is represented by a node in the graph, with two features per node: (i) the atomic species and (ii) the number of attached hydrogen atoms (one-hot vector). Edges are established between any 2 nodes within 3 covalent bonds of one another. Edge features consist solely of a one-hot vector indicating whether an edge represents a 3-hop, 2-hop, or 1-hop edge, where a 1-hop edge is equivalent to a covalent bond (*Figure* 1A).

**Fig. 1.**
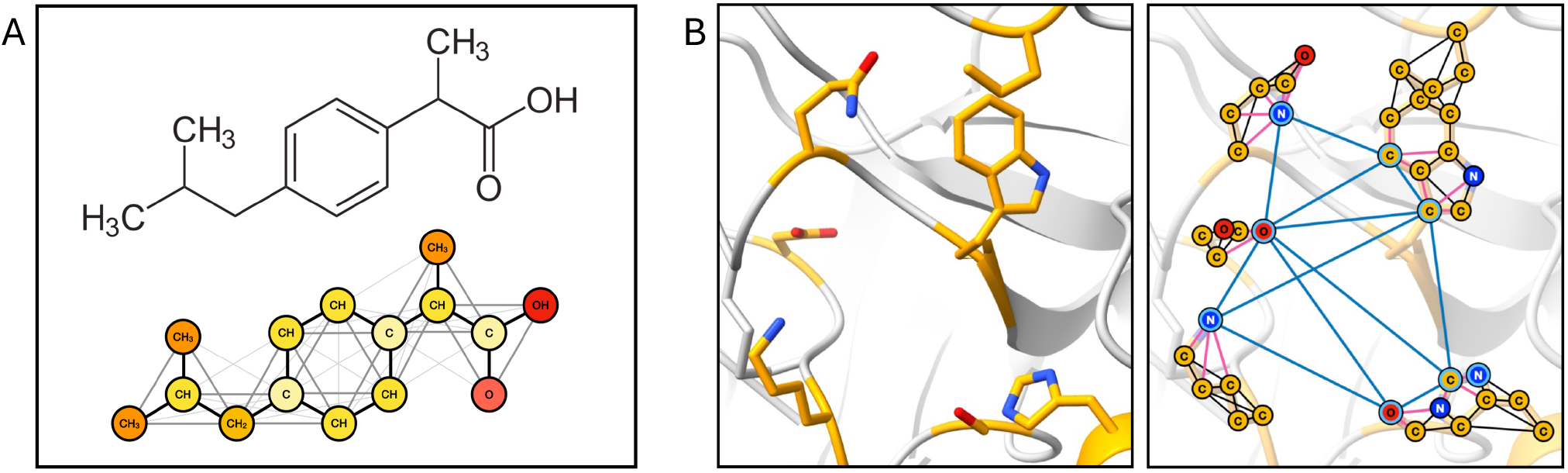
Graph Construction. In both compound graph and protein pocket graph, nodes represent heavy atoms. (A) In the compound graph, edges are created based on covalent bonds, representing one-hop, two-hop, and three-hop connections. (B) For a protein pocket graph: given a protein structure and a set of pocket surface atoms, edges are created in a neighborhood around pocket surface atoms. See text for details.

### Pocket Graph Construction

`Simpatico`’s algorithm requires specification of a target pocket. All protein structures used in training and evaluation include a ligand structure bound to the target pocket. Therefore, pocket specification begins by identifying all protein atoms within 6 Å of any ligand atom from the bound structure. This set of protein atoms is considered to make up the binding surface of the pocket, and thus referred to as the ‘pocket surface atoms’ **S**. This subset of the protein-atom point cloud is represented by the blue-outlined atoms in *Fig*. 1B.

All protein atoms that are not within 8 Å of a pocket surface atom are trimmed from the set of graph nodes. Edges are then established between any two atom nodes within 5 Å of one another. A virtual node is appended to the graph for each pocket surface atom, and given the same 3D coordinates as its corresponding protein atom. Directed edges are set from any protein atom within 5 Å to neighboring virtual nodes. Finally, edges are established between any two virtual nodes within 5 Å.

The object of this graph construction is to produce increasingly refined representations from the course atom level to that of the virtual nodes. In the supplementary material, we show that this approach confers a slight advantage over standard radius-based graph construction.

### GNN Architecture

Thus far, we have described the structure of the ligand and pocket atom graphs. These graphs are instantiated in two separate GNNs. All GNN model layers use Pytorch’s GATv2Conv, using multi-headed attention at each layer. The models are residually layered; each block consists of two GAT operations, the output of which is passed through a nonlinear SiLU layer and appended to the residual block input.

For pocket graph model edge features, distances between adjacent atom nodes are passed through a single low-dimensional linear layer with sigmoid activation. The output of this operation is used as the edge weight, which is the sole edge feature in the pocket GNN. Model details are provided in the supplementary materials.

### Training

Training resembles the precedent set by CLIP, which was designed to co-embed images with the correct descriptive text. In the case of `simpatico`, we seek to co-embed physico-chemically complementary pairs of protein and ligand atoms. Here, the standard of complementarity is observing both atoms in proximity (within 4 Å) in a resolved crystal structure.

For each step in training, a random batch ℙ𝕄 of protein-ligand graph pairs is retrieved. Any given pair *P*_*q*_*M*_*q*_ ∈ ℙ𝕄 contains some number of interacting protein-atom pairs such that *p*_*q,i*_ ∼ *m*_*q,j*_. Together, they represent a positive pair, comprising *p*_*q,i*_ as the anchor and *m*_*q,j*_ as the positive sample. The contrastive loss function is designed such that the loss is minimized when positive pairs are embedded closely in representation space.

For each positive pair, there is a corresponding selection of negative pairs, which should be placed far apart in embedding space. See below for `simpatico`’s process of selecting negatives during training. `Simpatico`’s model outputs *L*_2_-normalized embeddings of unit-length, so that the inner product between two embeddings directly corresponds to their euclidean distance. The loss produced for a single positive pair is then the cross entropy value calculated after the positive-pair and negative-pair inner products are concatenated and soft-maxed.

#### Intra-Molecular Negatives

Under a straightforward CLIP formulation, negative samples for an anchor protein atom could be sourced from random molecules from the batch. However, this strategy has a critical weakness: because the negatives are solely derived from random molecules, the model can learn to simply discriminate atoms based on their parent molecule, and subsequently collapse the atomic representations within a molecule to a single global embedding. While this is sufficient to minimize the contrastive loss function, it fails our objective of producing effective atom-specific representations.

To prevent representation collapse, we employ a notion of intra-molecular negatives: for a positive atom pair *p*_*q,i*_ ∼ *m*_*q,j*_, negatives are chosen from the same small molecule *m*_*q*_ (highlighted red in Figure 2). The risk of this strategy is that we inadvertently select false negatives – atoms that are not considered interaction partners based on the given structure, but are likely to experience periodic proximity in a real-world dynamic binding event (highlighted gray in Figure 2). An additional challenge is an atom that is so topologically proximal to the interacting ligand atom (1 or 2 covalent bonds away) that it is impossible for the model to adequately separate their representations.

**Fig. 2.**
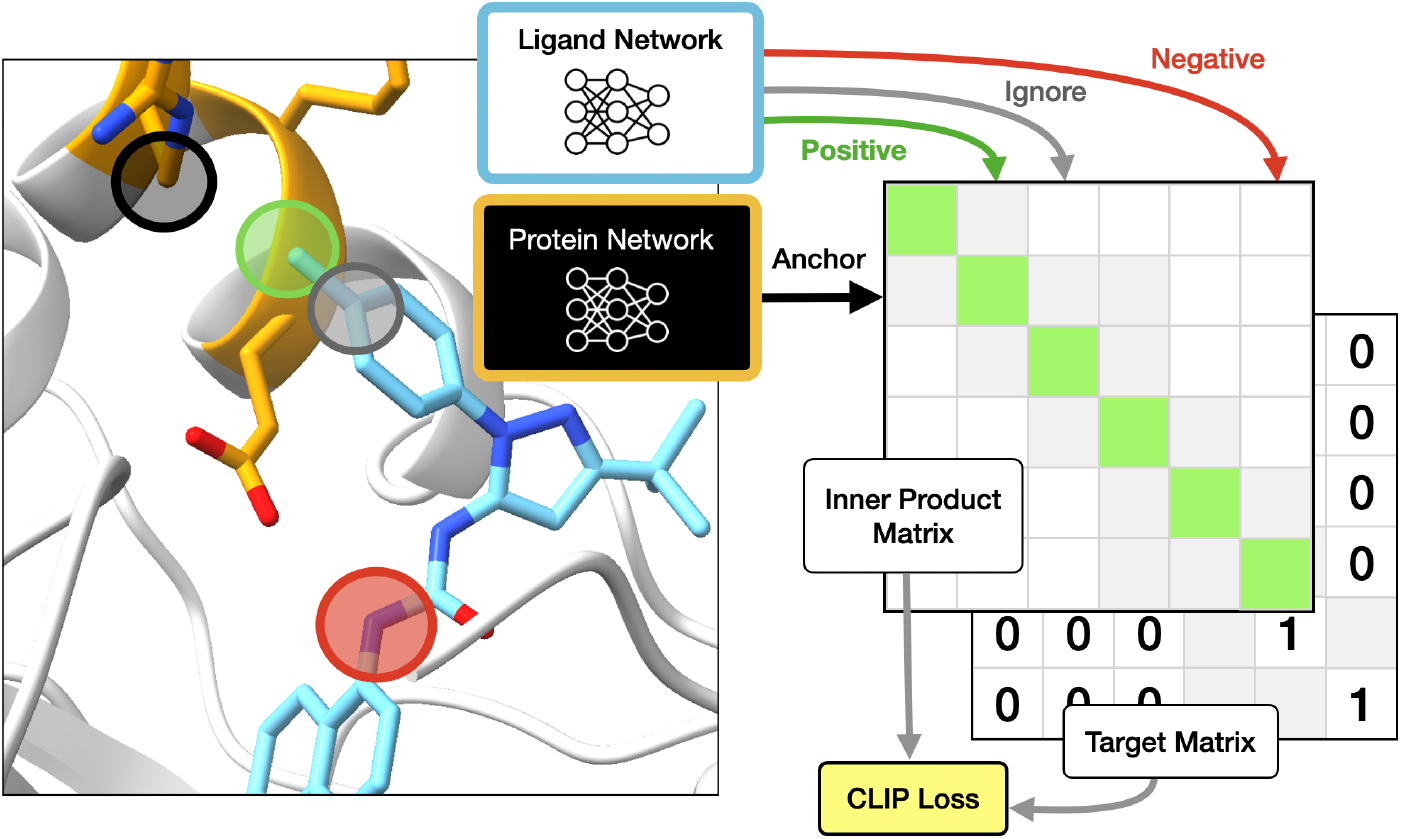
Intra-Molecular Loss. Following a modified CLIP loss framework: protein pockets (orange) and their corresponding bound ligands (light blue) are passed through separate GNN models to produce protein and ligand atom embeddings. The embedding values of highlighted circles on the left are indicated by colored arrows extending from the networks. For any two interacting protein-ligand atom pair, the protein atom (highlighted black) is used to produce the *anchor* embedding. The proximal ligand atom (highlighted green) is the source of the *positive* embedding. Atoms that are sufficiently distant from the interaction, though still belonging to the same small molecule (highlighted red) are marked as negatives. The CLIP loss function is optimized by maximizing the inner product between the anchor and positive embeddings (the positive inner product), while driving down the negative inner products. The inner product is ignored for the anchor and embeddings of atoms that are not interacting but are very close to interacting atoms (gray circle); these do not factor into the loss function. Cells corresponding to ignored inner products are shaded gray in the inner product and target matrices.

To minimize the selection of these false negatives, we establish a physical distance threshold of 6 Å. For any interacting protein atom *p*_*q,i*_ ∼ *m*_*q,j*_, all molecule atoms farther than 6 Å from *p*_*q,i*_ are considered as valid negatives.

#### Hard-Curriculum Negatives

Intra-molecular negatives encourage the model to produce atom-specific representations within a single molecule, but they do not require the model to integrate global structural context: A model trained with intra-molecular negatives alone can satisfy the contrastive objective by learning purely local chemical-group compatibility, without accounting for constraints imposed by the broader composition of the source molecule. For the models to learn an embedding strategy that accounts for these constraints, we should identify negatives with ostensible binding compatibility at the local level (i.e. traditionally compatible H-bonding partners) that we would nevertheless never see interacting in reality due to the incompatibility of their neighboring atoms. This requires selecting suitable negatives from separate molecules.

A naive selection strategy is to sample inter-molecule negatives uniformly at random from the training batch. However, in a batch of 16 protein-ligand pairs, most candidate negatives are too dissimilar from the positive to provide useful discriminative signal. We therefore construct an additional, larger pool of small molecules from which which negatives are drawn, and select from this pool using a curriculum that prioritizes negatives close to the model’s current decision boundary.

Specifically, for each anchor protein atom embedding 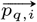, we compute the inner product between 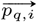 and the positive sample 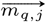, and between anchor 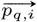 and every candidate negative atom in the pool. We then rank all atoms (positive plus candidate negatives) by inner product with the anchor, locate the positive’s position in this ranking, and sample negatives from a fixed-width window which overlaps that position. By construction, sampled negatives have inner products with the anchor that are similar to the positive’s, placing them near the model’s current decision boundary.

This is a form of curriculum learning [3]: as the model improves and the positive’s inner product with the anchor increases, the window shifts to harder negatives automatically, without an explicit difficulty schedule. See supplementary material for a technically detailed explanation of the curriculum learning procedure.

### Vector Databases

`Simpatico`’s retrieval step uses a vector database (FAISS [7]). After training, embeddings for every heavy atom of every small molecule in the screening library are written to the index. At screening time, each protein-atom embedding is used as a query, and FAISS’s fast nearest-neighbor search returns the *k* closest ligand-atom embeddings, i.e. the atoms most physicochemically compatible with that query. Queries from all atoms of a target pocket are issued in parallel, so the entire pocket is searched against the library in a single batched call. Query time grows sublinearly in index size, so the retrieval step for a pocket reduces to one batched lookup regardless of library size.

### Computing a score for a candidate pocket-ligand pair

In order to compute a scoring function (*P, M*) → ℝ^+^ for the binding potential between a protein pocket *P* and drug candidate *M*, `simpatico` employs a simple mechanism for accumulating molecule-level support from atom-level embeddings.

For a target pocket *P*, the vector-database retrieval step produces, for each 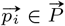, the set 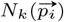 of nearest ligand-atom neightbors from 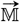.

An embedding similarity *s*(*p*_*i*_, *m*_*j*_) for each 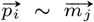 captured as described in the next paragraph. For each candidate ligand *M*_*n*_, the score *S*(*P, M*_*n*_) is computed as the sum of similarities of all identified neighbors 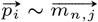:

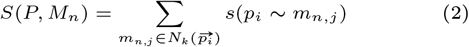

A measure of similarity *s*(*p*_*i*_, *m*_*j*_) for an atom pair is computed by calculating the Euclidean (L2) distance of the pair, and transforming this distance relative to (i) the greatest distance among all candidates pairs and (ii) a noise factor.

First, a maximum pairwise (Euclidean) distance is identified among a large sample of random *p*_*i*_, *m*_*j*_ pairs: 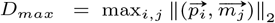. All pairwise distances are converted to similarities by subtracting the pair distance from 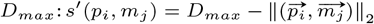. A result of this calculation is that the most distant ligand-atom pair will have a similarity of 0, and the pair with smallest distance will have a score close to *D*_*max*_.

These similarity scores are then adjusted in an attempt to remove the influence of random atom-pair matches on the accumulated score of a pocket-ligand pair. `Simpatico` computes the distribution of pairwise (Euclidean) distances between random atoms, and a noise threshold value t is identified such that 99% of all non-interacting pairs (*p*_*i*_ ≁ *m*_*j*_) have *s*′ (*p*_*i*_, *m*_*j*_) < *t*. This value is then subtracted from all *s*′ (*p*_*i*_, *m*_*j*_) values, with any resulting negative values clamped to zero: *s*(*p*_*i*_, *m*_*j*_) = max{0, *s*′(*p*_*i*_, *m*_*j*_) − *t*}.

## Results

### Enrichment Factor

To evaluate the efficacy of `simpatico` as a binding activity filter, we measure the extent to which it can filter away large amounts of a target database without filtering out the active molecules in that database. In practice, if `simpatico` can reliably produce small, strongly enriched samples from much larger datasets, it can enable an accurate overall workflow by limiting compute-heavy work to only the small enriched dataset. The *enrichment factor* of a sample is a measure of how many more-than-expected active molecules remain after down-sampling a database. Specifically: for a target protein, a compound dataset *S* is composed of actives *A* and negatives *N*, so *S* = *A* ⋃ *N*. `Simpatico` assigns scores to each compound, and the top *d*-percent scoring entries from *S* can be retained in a set called *S*_*d*_. Let *A*_*d*_ = {a|a ∈ *A* and *a* ∈ *S*_*d*_}. The enrichment factor of *S* at *d*% is:

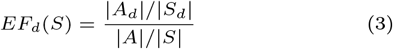

### Virtual screening experiments

The intended function of `simpatico` is to rapidly and accurately identify good candidate drugs for a target protein pocket. In general, this will be achieved by using `simpatico` to assign scores to compounds in a large database and to treat high-scoring compounds as good drug candidates. Note that `simpatico` does not produce a predicted docked structure; it simply assigns scores to good candidates, with the intention that some other (slower) tool will be used to perform docking on a candidate pool that is substantially enriched by `simpatico` for active molecules.

We evaluated `simpatico` on three primary benchmarking datasets: DEKOIS [2], DUD-E [18], and LIT-PCBA [26]. All three provide collections of proteins with a combination of known active compounds and a large number of challenging ‘decoy’ compounds that (a) are either known or expected to have no binding activity with the target protein and (b) are intended to be difficult to distinguish from actives by virtue of sharing similar properties with the actives. We also tested `simpatico` in a massively-high-throughput screening test involving a target database of 100 million candidates.

Different model weights were used per benchmark, each trained on the corresponding filtered subset of PDBBind described in Methods (Data). All reported values were obtained from the best performing weights observed over 100 epochs.

### Measuring efficacy with challenging decoy databases

#### DEKOIS results

The DEKOIS data set contains 81 protein targets. For each target, DEKOIS provides 40 known actives and 1200 property-matched decoys, yielding a total of 97,200 decoy molecules across the full set.

Table 1 shows the performance of `simpatico` relative to a variety of docking-based methods, presenting mean enrichment factor within the top 0.5%-scoring ligand candidates for each protein. These docking methods fall into one of two categories: classic energy-based or modern deep-learning based methods.

**Table 1.** Enrichment factors (EF_0.5%_) for multiple methods on DEKOIS dataset, consisting of 81 protein targets, each with 40 known actives paired with 1200 property-matched decoy, for a total of 3,240 actives and 97,200 decoys. Results for docking tools were taken from [5].

| Method | TANKBind | Vina | Gold | LeDock | Surflex-Dock | Glide SP | KarmaDock | SurfDock | **Simpatico** |
| --- | --- | --- | --- | --- | --- | --- | --- | --- | --- |
| $EF_{0.5\%}$ | 2.83 | 5.46 | 5.75 | 6.78 | 8.36 | 13.63 | 17.43 | 21.00 | **21.37** |

We highlight here that for this benchmark, `simpatico` was trained on a subset of PDBBind where proteins with 90% sequence identity to any target in the DEKOIS test set were removed. To our knowledge, no such data separation was performed for the deep-learning based docking methods, so that their results may benefit from some model leakage.

#### DUD-E and LIT-PCBA

In order to compare our results to several other methods, we performed screening on DUD-E and LIT-PCBA benchmarks. Table 2 presents the mean observed enrichment factors when evaluating the 1% top-scoring ligands for each protein. `Simpatico` enrichment factor values are much greater than classical docking approaches and also exceed state-of-art deep learning-based pose re-scoring such as Gnina. `Simpatico`’s run times are at least 25,000x faster than docking methods run on a single core. `Simpatico` also produces greater enrichment than DrugCLIP (note that both the `Simpatico` and DrugCLIP results are for models trained after removing training sequences with ≥ 90% identity to sequences in the test set).

**Table 2.**
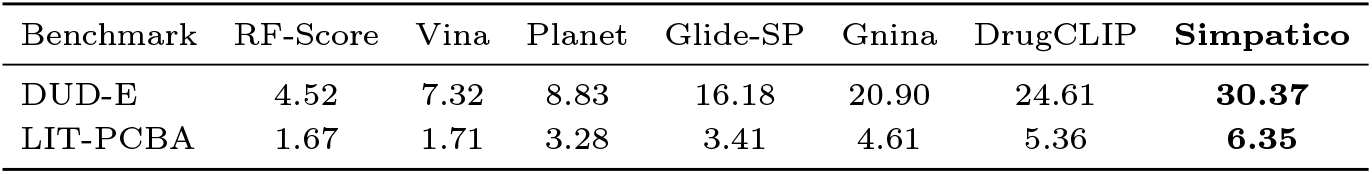
Enrichment factors (EF_1%_) on DUD-E and LIT-PCBA benchmarks. DUD-E has 102 proteins, 22,805 actives, and 1,144,300 decoys. LIT-PCBA has 15 proteins, 7,844 actives, and 407,381 decoys. EF values are taken from [14].

| Benchmark | RF-Score | Vina | Planet | Glide-SP | Gnina | DrugCLIP | **Simpatico** |
| --- | --- | --- | --- | --- | --- | --- | --- |
| DUD-E | 4.52 | 7.32 | 8.83 | 16.18 | 20.90 | 24.61 | **30.37** |
| LIT-PCBA | 1.67 | 1.71 | 3.28 | 3.41 | 4.61 | 5.36 | **6.35** |

**Table 3.** HTVS results: enrichment factor and total # actives retained among all 102×R top-scoring candidates.

| | 1%<br>($R=1$M) | 0.01%<br>($R=10$K) | 0.0001%<br>($R=104$) |
| --- | --- | --- | --- |
| EF (Enrichment Factor) | 57.3 | 2165.8 | 80,411.0 |
| # actives retained | 13,108 | 4646 | 1419 |

### High Throughput Virtual Screen

A significant strength of `simpatico` lies in its speed; its embedding retrieval strategy, leveraging a vector database for efficient neighbor search, allows it to avoid consideration of most candidates that have little chance of interacting with the protein-pocket target, enabling efficient search over massive chemical databases. To assess `simpatico`’s utility in for large-scale search, we performed a virtual screen using the proteins and associated ligands from DUD-E, mixed with 103,535,247 randomly selected compounds from the Enamine REAL collection of 13.6 billion compounds [9]. When actives from the DUD-E database are included, this results in a screening database of 103,558,052 small molecules.

`Simpatico` atomic embeddings for all actives and all sampled Enamine compounds were generated and sharded across ∼1000 separate files, each containing embeddings for ∼100,000 compounds (empirically, this was a number of ligands whose atom embeddings would fit in the 48GB RAM of our testing system’s NVIDIA L40S card).

Virtual screening was performed according to the scoring protocol described above. Values for *D*_*max*_ and noise threshold *t* were determined from the first shard of 100,000 embeddings, then used across all other shards. Each protein atom gathered the 2048 nearest ligand atoms in each shard, resulting in a total of ∼2 million nearest neighbor ligand atoms for each protein atom. Protein-compound scores *S*(*P, M*_*n*_) were computed for each compound M_*n*_ by summing over supporting atoms, as described in Methods.

We evaluated enrichment factors at down-sampling values of 1%, .01% and 0.0001%, corresponding to the top ∼1M, ∼10K, and ∼100 scoring molecules out of the ∼100M molecules screened respectively (for each target protein). Run time for search of the ∼100M ligands for matches to a single protein (AA2AR) required only ∼ 26 minutes when performed on a single L40S GPU; when all 102 proteins were searched as a batch, the highly parallel search completed in 5.65 hours. This compares favorably to the expected 300+ years of compute that would be required for such a screen performed with a docking tool that optimistically performed 1 docking operation per second.

These values reflect the runtimes required of the virtual screening search process, presuming that atom embeddings of the small molecule library have been pre-generated. In practice, generating embeddings of the ∼ 100M small molecules required ∼10 hours on an NVIDIA V100S-32GB GPU. Note that in `simpatico`’s expected use-case, this process needs only be run once, after which the user may perform rapid screenings on any number of arbitrary protein targets.

Of the 22,805 DUD-E actives, 20% (4646) were retained in the top 10,378 scoring compounds for each protein (0.01% downsampling). Even when sampling only the top 0.0001% scoring compounds (104 for each protein), 6% of all actives were retained, with a mean of 14 actives retained per protein. In one example, of the 100 actives corresponding to protein target NOS1, 90 were among the 104 top-scoring of all ∼100M compounds.

### Ablation Tests on the Choice of Negatives

To assess the contribution of each of `simpatico`’s two negative-sampling strategies (intra-molecular and hard-curriculum, both described in Methods), we trained the model twice with one of the two negative sources disabled in each run. The resulting performance on the DUD-E dataset is available in Table 4. Clearly, the use of both sources of negatives is crucial to `simpatico`’s discriminatory power.

**Table 4.**
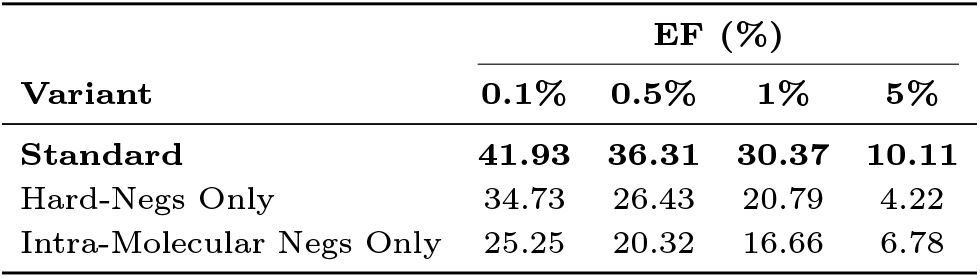
Ablation study results for Enrichment Factor (EF) percentages.

| Variant | EF (%) |  |  |  |
| --- | --- | --- | --- | --- |
|  | 0.1% | 0.5% | 1% | 5% |
| **Standard** | **41.93** | **36.31** | **30.37** | **10.11** |
| Hard-Negs Only | 34.73 | 26.43 | 20.79 | 4.22 |
| Intra-Molecular Negs Only | 25.25 | 20.32 | 16.66 | 6.78 |

### T-SNE Analysis of Pocket-Ligand Pairs

We performed T-SNE clustering of the embeddings of atoms belonging to a collection of 10 different protein-ligand complexes (Figure 3, Left). Because our training objective seeks to co-locate interacting pairs of atoms, we should expect to see ligand and protein atoms clustered according to their interactions, not just their parent complex. To show that this is accomplished by our embedding strategy, we zoom into a particular cluster (pink box) to inspect the individual embeddings, as seen in Figure 3. Here, protein atoms are once again represented by circles and ligand atoms by triangles. To generate the color, we took the coordinates of the most physically-distant pair of ligand atoms in the cluster, designated poles 1 and 2, and produced a value for each a from the following function:

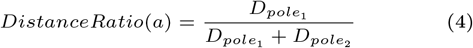

Where *D*_*pole*_ is the distance in angstroms from atom a to pole *i*. The effect of this function is that atoms very near pole 1 receive a value very near 0 (blue), and atoms very near pole 2 receive a score very near 1 (red), while atoms that are not especially close to either point will tend towards intermediate values (gray). From Figure 3, we observe that ligand and protein atoms physically proximal to pole 1 also tend to cluster together in embedding space, while ligand and protein atoms observed near pole 2 are found near one another in embedding space. The embedding space successfully embeds atoms according to their interactions, and not just by their target protein. Other clusters may be viewed in the supplementary material.

**Fig. 3.**
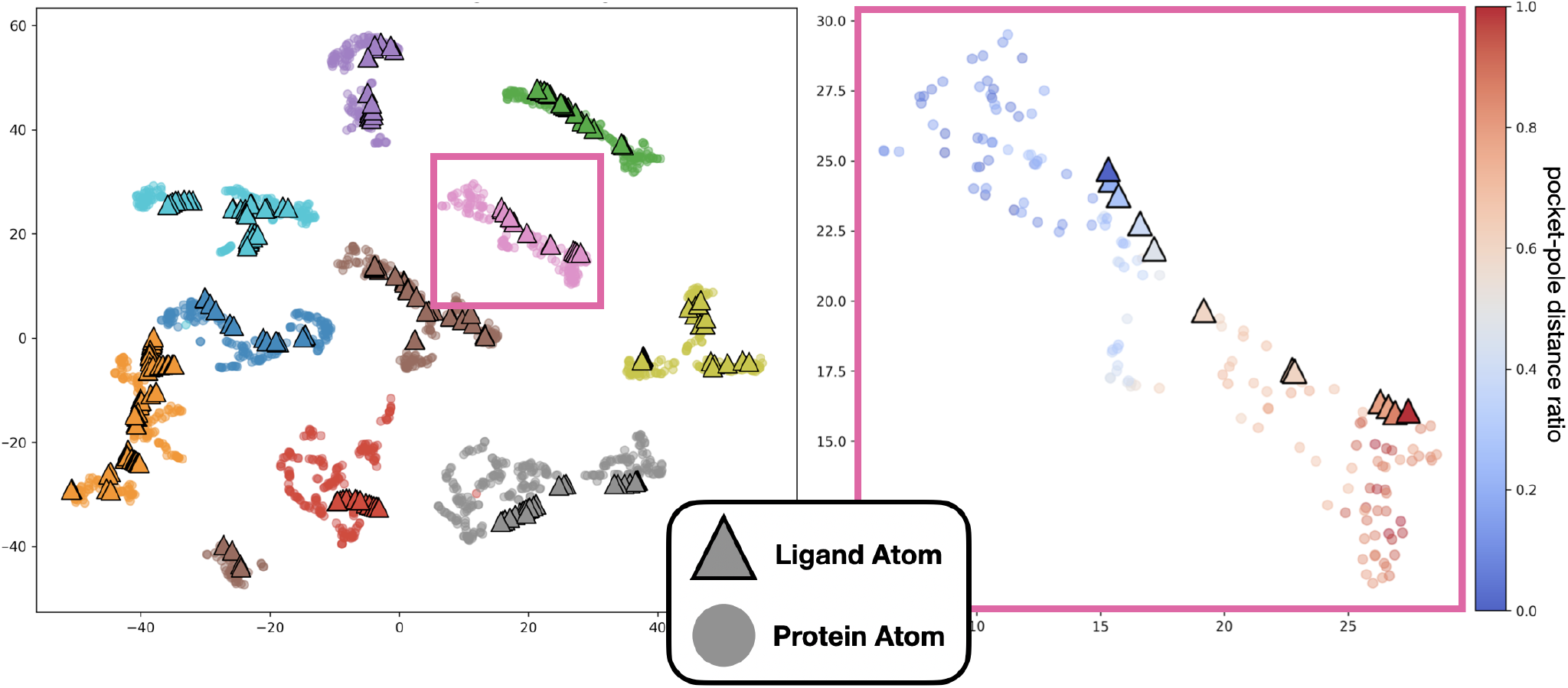
(Left) T-SNE visualization of atomic embeddings from ten different protein-ligand pairs produced by the same model trained on different loss functions. Protein atom embeddings are indicated by a circle and ligand atom embeddings by a triangle, both colored by their protein-ligand complex, with non-interacting (i.e. greater than 6 Å from any ligand atom) protein atoms filtered out. We observe good clustering of protein pocket atoms and their binding partners. (Right) We zoom into the cluster bounded by the pink box and color-code the ligands and atoms according to their proximity to the two most distant points on the ligand structure. This shows good agreement in embedding value between atom neighbors.

### Per-atom matching outperforms whole-molecule pooling

`Simpatico`’s advantage over methods like DrugCLIP could plausibly be due to incidental differences in training data or architecture rather than to per-atom matching itself. To isolate the contribution of the per-atom approach, we evaluated a family of models that are nearly identical to Simpatico except for the addition of a final aggregation step that collapses the per-atom embeddings into a single molecule-level (or pocket-level) representation, used for retrieval in the same CLIP-style manner as DrugCLIP. We focus on attentional aggregation, which gives the model the opportunity to weight chemically important atoms more heavily; for completeness we also report a parameter-free mean-pooling variant of each model. The hard-negative curriculum from atom-level training is preserved throughout. We evaluate four training regimes (Table 5):

**Table 5.** Pooling baselines compared with `simpatico` (per-atom matching) on DUD-E, evaluated by enrichment factor at 1% (EF_1%_). Variants differ in training loss and embedding dimension; the 512-dimensional configuration was applied only to the best-performing 64-dimensional variant (Joint Loss). In the atomic-loss column, P denotes atomic loss applied during pretraining only (the encoder is then frozen while the aggregator is trained with a molecular loss), whereas ✓ denotes atomic loss active throughout training.

| Variant | Mol. loss | Atomic loss | Dim. | Attention | Mean |
| --- | --- | --- | --- | --- | --- |
| Frozen Atomic Encoder | ✓ | P | 64 | 18.45 | 20.16 |
| Molecular Only | ✓ | | 64 | 18.82 | 20.18 |
| Joint Loss | ✓ | ✓ | 64 | 19.84 | 21.81 |
| Joint Loss (512) | ✓ | ✓ | 512 | 23.47 | 21.56 |
| **simpatico (per-atom)** | | ✓ | 64 | **30.37** | |

#### Frozen Atomic Encoder

The encoder is trained with the per-atom interaction objective as in standard `simpatico`, then frozen; only the attention layer is trained, supervised by a contrastive molecule-level loss. For the mean-pooling variant no training is needed: the molecule embedding is simply the average of the frozen atom embeddings.

#### Molecular Only

The encoder and attention layer are trained end-to-end from scratch using only the molecule-level loss; the per-atom interaction objective is dropped entirely.

#### Joint Loss

Same as Molecular Only, but the loss adds Simpatico’s per-atom interaction objective to the molecule-level objective, on the intuition that per-atom supervision may keep interaction-level signal from collapsing into the pooled representation.

#### Joint Loss (512)

A repeat of Joint Loss with the output dimension widened from 64 to 512, matching the dimensionality used by DrugCLIP and UniMol. This isolates whether 64 dimensions is simply insufficient capacity for whole-molecule retrieval.

None of the pooled variants approach `simpatico`’s enrichment EF_1%_ of 30.37. Neither widening the embedding nor adding the per-atom auxiliary loss is enough to match `simpatico`: the auxiliary loss yields only modest gains in either pooling style, and even at 512 dimensions (matching DrugCLIP and UniMol) the best pooled variant reaches only 23.47. The mean-pooling variants are within two points of their attention counterparts, and in the smaller-dimensional regimes actually edge them out, suggesting the attention layer is not learning to identify binding-relevant information that pooling otherwise discards. Together, these results indicate that the gap between Simpatico and pooled-embedding retrieval is not an artifact of training data, training objective, or capacity, and instead reflects information that any single-vector summary necessarily loses.

## Discussion

We have introduced a novel strategy for identifying promising candidates for protein-ligand binding activity. Classical docking approaches and newer deep learning-based methods require a great deal of computational power, restricting their application on billion-scale search. However, a new paradigm has emerged that leverages advances in representation learning to make this process drastically more efficient, by retrieving good ligand candidates from a massive database based on their locations in embedding space. `Simpatico` approaches this problem by leveraging a novel loss function to co-locate the embeddings of atoms of small-molecules and protein-pocket residues according to their interaction potential. The binding potential between a protein pocket and small molecule then becomes a function of the individual interactions we expect to observe during binding. `Simpatico` demonstrates substantial gains in both speed and accuracy over traditional methods, as well as state-of-art performance in the dense-retrieval approach to virtual screening.

In light of `simpatico`’s superior performance, it is important to highlight several aspects of our approach that contrast with other dense-retrieval methods (as exemplified by the recently-released DrugCLIP method).

### Why per-atom matching is effective

The first, and perhaps most crucial innovation of `simpatico` is performing retrieval on the basis of *individual atomic interactions* rather than whole-molecule representations. We demonstrated this by minimally adjusting model architectures to produce whole-molecule embeddings, and showed that these do not achieve the same screening performance as the atomic-embedding approach. Figure 3 offers some insight into why this may be: compatible protein-ligand atom embeddings cluster with their binding partner in embedding space, and because each ligand atom can be independently retrieved as a near-neighbor of a complementary protein atom, each candidate ligand has multiple opportunities to be retrieved during search. In this sense, the aggregation of individual atom scores may be seen as a collection of “votes” on how likely the parent molecule is to bind.

Figure 4 reinforces this view. To produce the figure, the atomic embedding values in a single protein-ligand complex were reduced to 3 dimensions via a standard PCA reduction. Subsequently, each atom was assigned the RGB color corresponding to its 3-dimensional value after normalization. When these embedding-derived color values are applied to a 3D representation of the ground-truth complex structure, we see considerable agreement between ligand atom embeddings and their protein atom neighbors. When the protein pocket atoms are used to query ligand embedding space for ligand atom partners, the source ligand is likely to accumulate a strong score.

**Fig. 4.**
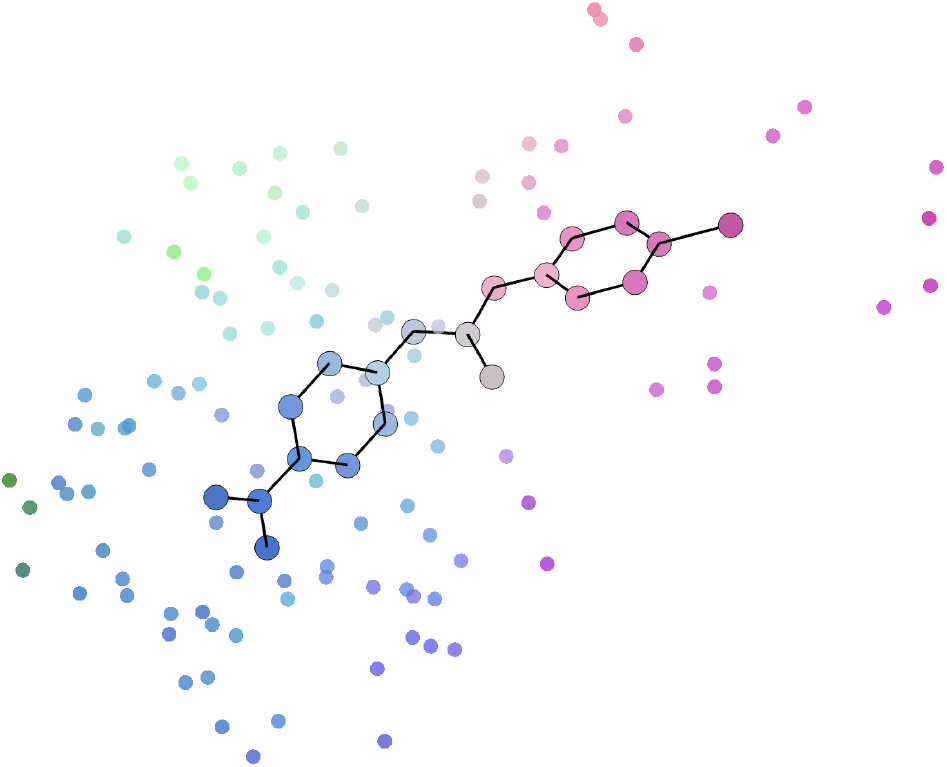
3D representation of molecule GP6 in complex with Beta-Trypsin binding (PDB id 1BJU). Protein pocket atoms are depicted without covalent bonds. Protein and ligand atomic embeddings values were reduced to the top 3 primary components and normalized along their respective dimensions. Atoms were then colored with the RGB value corresponding to this final 3-dimensional representation. The operation shows agreement in embedding space between protein and ligand atoms that are approximate in 3D space, and thus chemically complementary.

### Better aggregation could substantially improve performance

Because `simpatico`’s pipeline cleanly factors into representation and aggregation, the upper bound on aggregation performance for a fixed representation is straightforward to compute by calculating the enrichment factor that would have been achieved if the aggregation step had performed perfectly, ranking all actives higher than any decoy. Table 6a compares such an oracle against our current aggregation step.

**Table 6.** Oracle results on three benchmarks (DUD-E and LIT-PCBA).

| (a) DUD-E |  |  |  |
| --- | --- | --- | --- |
| Sampling Ratio | 0.1% | 0.5% | 1% |
| Standard | 41.93 | 36.31 | 30.37 |
| Oracle | 63.08 | 62.99 | 59.78 |

| (b) LIT-PCBA |  |  |  |
| --- | --- | --- | --- |
| Sampling Ratio | 0.1% | 0.5% | 1% |
| Standard | 23.38 | 8.89 | 6.35 |
| Oracle | 151.08 | 45.21 | 24.37 |

The same pattern holds, more dramatically, on LIT-PCBA (Table 6b): the oracle reaches an EF_0.1%_ of 151.08 against our current 23.38, roughly four times the proportional gap observed on DUD-E. This suggests that on harder benchmarks the aggregation step, rather than the per-atom representation itself, is the dominant bottleneck, and that improvements to aggregation may yield the largest gains precisely where current performance is most limited.

While other methods that provide per-molecule embeddings can only be improved by updating their representation scheme, `simpatico` can in principle be significantly improved through this or by developing new aggregation strategies. The performance gap between the current aggregation function and that of the perfectly performing oracle represents a potential avenue for improving our method overall.

### Structural information enables data-efficient training

`Simpatico` was trained from scratch, starting from one-hot atom-feature encodings rather than pre-trained embeddings from models such as UniMol or PLMs. The physical-interaction-based learning objective requires 3D structural training data, which is much less abundant than general protein-ligand binding data (PDBBind has ∼20K complexes, while BioLip contains hundreds of thousands of binding records). Further, removing proteins highly similar to test-set targets from the training data reduced the available data by as much as 20% for the DUD-E dataset.

Despite these obstacles in the availability of data, `simpatico` still outperforms current deep-learning-based docking and dense-retrieval virtual screening methods. This suggests that structural data is vastly more information-dense than binary protein-ligand binding labels combined with generic pre-trained embeddings for proteins and small molecules.

### Score accumulation

Our “scoring oracle” demonstrates a substantial gap between our current aggregation and the theoretical best, with the gap widening on the harder LIT-PCBA benchmark, suggesting aggregation is most limiting where the representation problem is hardest. The current score accumulation method is admittedly naive, and we expect more advanced alternatives to yield substantial improvements in screening. For example, accumulation methods that account for the ligand conformation implied by the atom-embedding pairings should help reject hits with strong individual interactions but poor overall fit for the pocket.

### Future work

We anticipate that large advances will result from improved models, better training data (for example using [28], or expanding to include molecular dynamics data from [22]), and schemes for accumulating support from atom pairs that improve on the simplistic approach currently employed in `simpatico`. We also note that we see promising improvements in both speed and accuracy in early tests in which we employ an HNSW database such as CAGRA [19] for vector near neighbor search.

One shortcoming of per-atom representation is that it creates the need to store many more embeddings for each ligand, creating a challenge where speed and storage are concerns. Though we have shown that the complete collapse of molecular structures into single representations significantly compromises performance, it will be worthwhile to explore methods for reducing the data requirements of storing atom representations of massive small-molecule libraries. There are many avenues that could aid in this endeavor, including dimension reduction techniques, pooling strategies, and vector quantization.

## Author contributions statement

J.G. and T.J.W. conceived the model design, training schemes, and experiment(s); J.G. implemented the software, performed training, conducted the experiment(s), and analyzed the results. J.G. and T.J.W. wrote and reviewed the manuscript.

## Acknowledgments

We thank Daniel Olson and Hoi Kei (Patricia) Woo for thoughtful feedback and suggestions throughout the course of development of this work. We are grateful for the high performance computing (HPC) resources supported by the University of Arizona TRIF, UITS, and Research, Innovation, and Impact (RII) and maintained by the UArizona Research Technologies department. Compute was also supported through Jetstream2 g3xl compute nodes provided by allocation CIS240916 from the Advanced Cyberinfrastructure Coordination Ecosystem: Services & Support (ACCESS) program, which is supported by National Science Foundation grants #2138259, #2138286, #2138307, #2137603, and #2138296. This work was supported by NIH NIGMS R01GM132600 and T32GM132008, an NIH PATHWAYS fellowship at Rocky Mountain Laboratories through NIAID Bioinformatics and Computational Bioscienc es Branch, DOE BER DE-SC0021216, and the DOE PerCon SFA, Secure Biosystems Design Project.

