## supplementary material for "Simpatico: accurate and ultra-fast virtual drug screening with atomic embeddings"

### Supplementary Materials

##### *Description of benchmark data sets (DEKOIS, DUD-E, and LIT-PCBA)*

The DEKOIS [2] data set contains 81 protein targets. For each target, DEKOIS provides 40 known actives paired with 1200 property-matched decoys, yielding a total of roughly 97,200 decoy molecules across the full set. DUD-E [18] includes 102 protein targets, with a total of 22,805 experimentally confirmed actives (an average of 224 actives per target). For each active ligand, 50 decoys are generated (similar in physico-chemical properties but dissimilar in topology), giving on the order of 1.14 million decoy compounds in total. LIT-PCBA [26] includes 15 distinct protein targets, each with experimentally confirmed actives and inactives. It contains a total of 7,761 true actives and 382,674 confirmed inactive compounds.

##### *Pocket graph construction: hierarchical vs simple radius-graph*

A simple method for producing a geometric representation of the protein pocket is to assign distance-weighted edges between any two atom nodes within some threshold distance of each other. We developed a slightly more complex scheme: a virtual node is established at the location of any protein atom within 6 Å of a template ligand coordinate. The graph is then constructed such that information flows from a simple radius-based graph of the pocket to the virtual nodes. Ultimately, the final embedding value of these virtual nodes are used as the target queries in the VS search process.
Below, we provide plot S1 to compare the test values from both versions over 100 epochs of training, where evaluation is performed on the DUD-E dataset every 5 epochs. For both versions, a threshold of 5 Å is set for all radius-based edge generation. Because the hierarchical version appears to confer a slight advantage, we elected to use this scheme for pocket graph construction throughout.

##### *Protein Encoder*

The `ProteinEncoder` generates 64-dimensional embeddings for protein pocket surface atoms using a hybrid graph of physical atoms and virtual nodes, processed through a GATv2-based residual architecture.

- **Input:** Each node has a 59-dimensional input feature vector, corresponding to a heavy (non-hydrogen) atom of the protein structure. Input vector indices correspond to the following features:
  - $[0 - 37]$: One-hot indicator of atom name.
  - $[38 - 58]$: One-hot indicator of parent residue.

  Atom features are projected to a 256-dimensional space (`hidden_dim` = 64, `heads` = 4).
- **Graph Construction:** The architecture utilizes three types of directed edges constructed via a radius-based search ($r = 5$, max $k = 32$):
  - *Atom–Atom:* Connections between physical atoms within the pocket.
  - *Atom–Virtual:* Bipartite connections between physical atoms and virtual nodes.
  - *Virtual–Virtual:* Connections between virtual nodes.

  Edge weights are generated by a `PositionalEdgeGenerator`, which applies a 2-layer MLP with Sigmoid activation to inter-node distances to bound weights between 0 and 1. During training, DropEdge regularization ($p = 0.2$) is applied to the global edge index to prevent over-fitting.
- **Graph Layers:** The network consists of 3 residual blocks (`blocks` = 3), each containing 2 `ResLayer` modules (`block_depth` = 2). Each layer follows a pre-activation residual structure:
  1. Layer Normalization of the input features.
  2. GATv2 convolution operating on 256-dimensional features with 4 attention heads.
  3. SiLU activation.
  4. Dropout ($p = 0.1$).

  The output of each residual block is concatenated with the initial input representation, preserving multi-scale feature information.
- **Output:** The resulting 1024-dimensional concatenated representation (derived from $3 + 1$ stages) is passed through an output MLP consisting of Layer Normalization, a linear layer with ReLU activation, and Dropout ($p = 0.1$). The final 64-dimensional virtual-node embeddings (`out_dim` = 64) are $L_2$ normalized and returned as output. These virtual node embeddings are intended for use as queries in the downstream virtual screening search.

*Small Molecule Encoder*

The `MolEncoder` produces 64-dimensional atom embeddings using a deep residual GATv2 architecture.

- **Input:** Each heavy atom in the small molecule is represented by a 33-dimensional feature vector. Input vector indices correspond to:
  - $[0-28]$: One-hot indicator of element name (RDKit conventions).
  - $[29-32]$: One-hot indicator of the number of adjacent hydrogen atoms.

  Atom features are projected into a 256-dimensional space (`hidden_dim` $= 64$, `heads` $= 4$) via an initial linear layer.
- **Edge Construction & Attributes:** The graph utilizes a topological edge construction where edges are established between atoms within 3 covalent bonds. Each edge is associated with a 3-dimensional one-hot vector encoding the topological distance (bond hops).
- **Graph Layers:** The encoder employs 6 residual blocks (`blocks` $= 6$), each containing 2 `ResLayer` modules (`block_depth` $= 2$). The layers implement a pre-activation residual structure to stabilize deep training:
  1. Layer Normalization.
  2. GATv2 convolution operating on 256-dimensional features with 4 attention heads and 3-dimensional edge attributes.
  3. SiLU activation.
  4. Dropout ($p = 0.1$).

  The outputs from every residual block are stored and concatenated with the initial projection, preserving multi-scale structural information across 1792 total dimensions.
- **Output:** The concatenated representation is passed through a refined output projection sequence:
  1. Layer Normalization of the 1792-dimensional concatenated vector.
  2. Linear layer projecting to a 256-dimensional bottleneck.
  3. SiLU activation and Dropout ($p = 0.1$).
  4. Final linear layer producing 64-dimensional embeddings (`out_dim` $= 64$).

  All final embeddings are $L_2$ normalized to unit length.

#### Hard-Curriculum Learning

Hard negatives were selectively mined from a separate, larger batch of ligand atoms. The basic idea is to use negative pairs with similar inner-products to their corresponding anchor-positive pairs. This way, the model learns to iteratively refine its representations to gradually update the decision boundary, without wasting resources on negatives that are either trivially distinguishable or too similar to provide a useful discriminatory signal.

For each training batch, a separate random batch of ligands is retrieved, separate from the ligands in the protein-ligand pairs, and passed through the small-molecule encoder to produce batch of negative atom embeddings.

The inner product between all anchor atoms and all negative atom embeddings is evaluated. Each row of anchor-negative inner products is then sorted, so that the anchor-positive inner product can be ranked among them. At large batch sizes, straightforward sorting of so many rows produced GPU memory failures. Therefore, we employ Pytorch's `torch.topk` to obtain and sort only the greatest 2048 inner product values per row.

In early training, with sufficiently large negative-ligand batches, the anchor-positive inner product will not rank among the 2048 highest inner products. Therefore, we begin with a very small batch size of 32 ligands. With an average of $\sim$30 atoms per small molecule, a batch size of 32 ligands will on average produce $\sim$960 small atom embeddings.

Over training, performance will improve, corresponding to higher ranking anchor-positive inner products. We may therefore gradually increase the size of the hard-negative pool as the average positive inner product ranking improves. In practice, we grow the hard negative batch by 50% if the average positive inner product rank remains among the 15% highest ranking inner products for 100 training batches, with a maximum hard negative batch size of 512.

Each anchor positive pair will have its own rank among the sorted negative inner products. The selection of negatives for each is drawn from window of length 100 over the sorted list, that always includes the positive inner product rank.

For each sample, the location of the window is based on the relative rank such that the ratio of better-scoring to worse scoring negatives is precisely equal to the relative rank. If an anchor-positive IP is ranked 512 among the 2048 sorted values, the window is situated such that 1/4 of the negatives rank better than then positive. This is implemented simply to handle cases where the window would extend outside of the sorted list (if rank=1, the window extends from index 0 to 99, and if ranked 2048, the window will extend from index 1947 to 2047).

In cases where the anchor-positive IP is not among the top 2048, it is simply given the last-ranking value of 2048.

#### Protein-Ligand Proximity Plots

In Figure 3, we showed how clustering occurs at the level of interacting atoms, not just pocket and ligand representations in general. Figure **??** shows clustering between the 9 remaining pocket-ligand pairs shown in Figure 3.

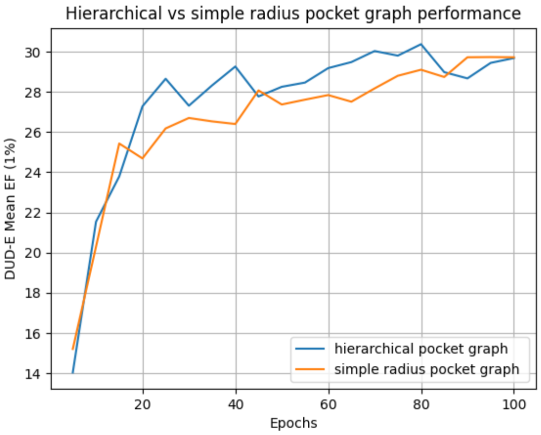


**Fig. S1.** Comparison of pocket graph constructions. DUD-E $EF_{1\%}$ over 100 training epochs (evaluated every 5 epochs) for the simple radius-based pocket graph and the hierarchical virtual-node graph described in the text. Both use a 5 Å threshold for radius-based edges. The hierarchical scheme yields a small but consistent advantage.

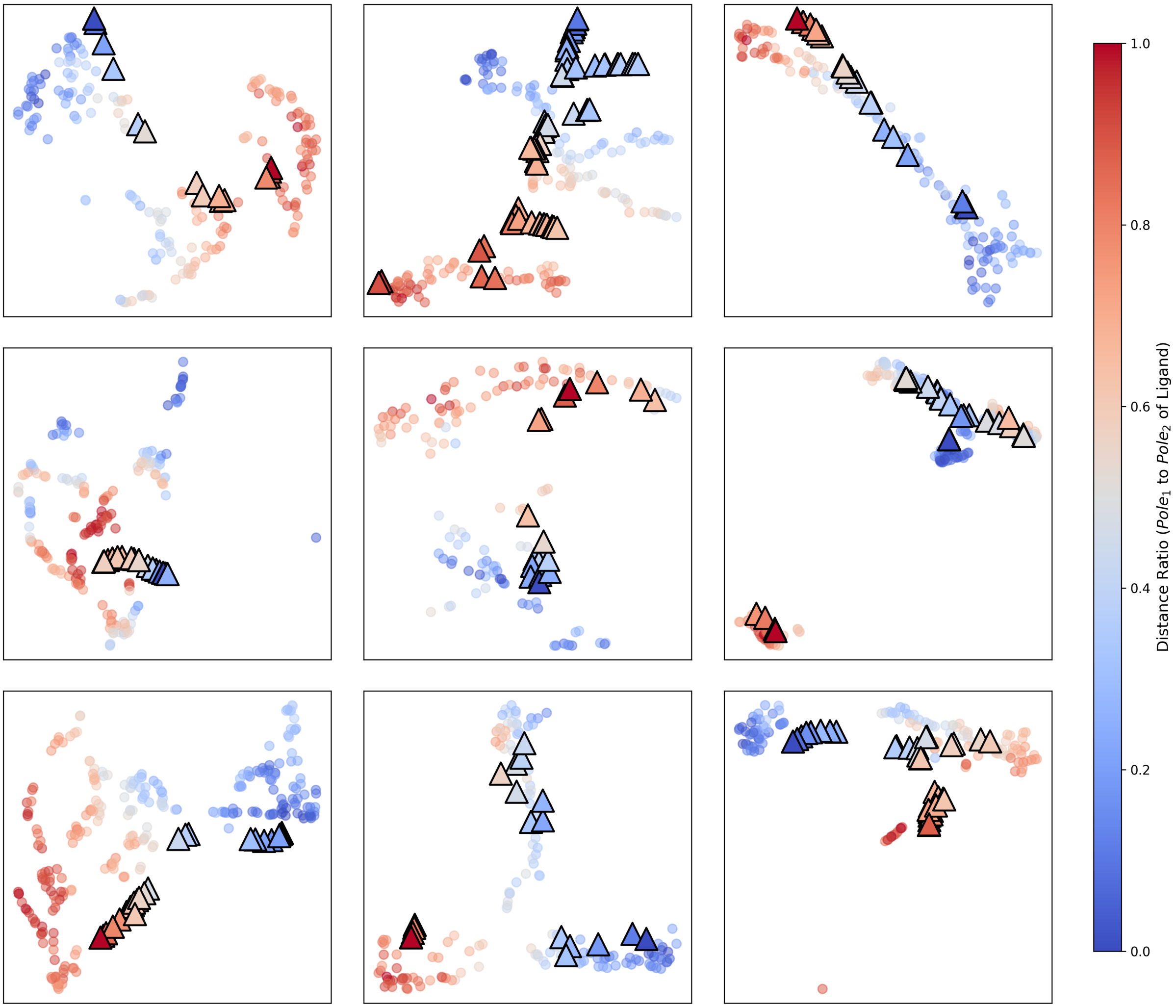


**Fig. S2.** T-SNE visualization of atomic embeddings for the remaining 9 protein-ligand pairs from Figure 3. Protein atom embeddings are shown as circles and ligand atom embeddings as triangles, colored by parent complex; non-interacting protein atoms (>6 Å from any ligand atom) are filtered out. As in Figure 3, clustering occurs between interacting protein and ligand atoms rather than only by parent complex.
